# Environmental variation mediates the evolution of anticipatory parental effects

**DOI:** 10.1101/606103

**Authors:** Martin I. Lind, Martyna K. Zwoinska, Johan Andersson, Hanne Carlsson, Therese Krieg, Tuuli Larva, Alexei A. Maklakov

**Affiliations:** Animal Ecology, Department of Ecology and Genetics, Uppsala University, 752 36 Uppsala, Sweden; School of Biological Sciences, University of East Anglia, Norwich Research Park, Norwich NR4 7TJ, UK

**Keywords:** *Caenorhabditis*, Environmental heterogeneity, Maternal effects, Reproduction, Temperature, Transgenerational plasticity

## Abstract

Theory maintains that when future environment is predictable, parents should adjust the phenotype of their offspring to match the anticipated environment. The plausibility of positive anticipatory parental effects is hotly debated and the experimental evidence for the evolution of such effects is currently lacking. We experimentally investigated the evolution of anticipatory maternal effects in a range of environments that differ drastically in how predictable they are. Populations of the nematode *Caenorhabditis remanei*, adapted to 20°C, were exposed to a novel temperature (25°C) for 30 generations with either positive or zero correlation between parent and offspring environment. We found that populations evolving in novel environments that were predictable across generations evolved a positive anticipatory maternal effect, since they required maternal exposure to 25°C to achieve maximum reproduction in that temperature. In contrast, populations evolving under zero environmental correlation had lost this anticipatory maternal effect. Similar but weaker patterns were found if instead rate-sensitive population growth was used as a fitness measure. These findings demonstrate that anticipatory parental effects evolve in response to environmental change so that ill-fitting parental effects can be rapidly lost. Evolution of positive anticipatory parental effects can aid population viability in rapidly changing but predictable environments.

**Impact summary:** Parents can help their offspring by adjusting offspring’s phenotype to match their environment. Such anticipatory parental effects would be beneficial, but only if parents can accurately predict the environment in which their offspring will develop and live. The plausibility of such anticipatory parental effects is hotly debated. The theory is clear that the predictability of the environment should play a defining role. We used an “experimental evolution” approach in a fast reproducing nematode worm *Caenorhabditis remanei* to tackle this question and follow the evolution of parental effects in different environments in real-time. We found that populations evolving in a novel but predictable environment indeed had anticipatory parental effects that increased fitness of their offspring in that environment. In contrast, when evolving in an unpredictable environment where such parental effects would be disadvantageous, the parental effect was rapidly lost in evolution. Our novel experimental environments were constructed by exposing worms to increased temperature. Anticipatory parental effects play an important role in adaptation to novel environments and will affect the viability of populations under climate heating.

## Introduction

The role of environmental variation in the adaptive expression of phenotypes has gathered considerable interest (Chevin et al. 2010; Hollander et al. 2015; Donelson et al. 2017; Lind and Spagopoulou 2018). Not only is heterogeneity common, it is also predicted to change evolutionary outcomes. While stable environments generally should select for genetic specialisation, environmental heterogeneity can select for environmental input on this process. Variable environments is expected to select for phenotypic plasticity (if environmental cues are reliable) or bet-hedging (if the cues are not reliable) (Moran 1992; Simons 2011). However, a developing organism may not be able to acquire and/or interpret the environmental cues itself; therefore, the parental environment can also function as a developmental cue (Marshall and Uller 2007; Leimar and McNamara 2015). Consequently, recent theory maintains that high environmental correlation results in predictability between generations, which can select for adaptive parental effects and/or epigenetic inheritance of an environmentally induced phenotype (Lachmann and Jablonka 1996; Bonduriansky and Day 2009; Rivoire and Leibler 2014; Kuijper and Hoyle 2015; Leimar and McNamara 2015; Uller et al. 2015; Proulx et al. 2017; Dury and Wade 2020), mechanisms collectively referred to as inter- or transgenerational plasticity (Perez and Lehner 2019). The sign of the environmental correlation is expected to result in a similar sign of the parental effect, so that parents can prepare their offspring for the same environment as they are themselves experiencing (positive correlation), the opposite environment (negative correlation) or does not influence the phenotype of their offspring (zero correlation between parent and offspring environments). The exception is constant (highly predictable) environments, where genetic specialisation is predicted to evolve (Leimar and McNamara 2015) together with a negative trans-generational effect in order to reduce phenotypic variance (Hoyle and Ezard 2012; Kuijper and Hoyle 2015).

However, while the theory is well developed, the empirical evidence is mixed. In its most basic form, the theory of anticipatory parental effects predicts that offspring should have higher performance if parental and offspring environments are matching (Marshall and Uller 2007; Burgess and Marshall 2014). There are some striking examples of this (e.g. Gustafsson et al. 2005; Galloway and Etterson 2007; Jensen et al. 2014; Kishimoto et al. 2017; Ryu et al. 2018; Toyota et al. 2019), but across studies there is only weak support for this prediction in natural systems, and the effects are small compared to the direct effects of offspring environment (reviewed in Uller et al. 2013). Since environmental predictability across generations is seldom quantified, one reason for the scarcity of clear examples of adaptive parental effects is that environments may not often be correlated across generations, and, therefore, provide little opportunity for selection for such anticipatory effects (Burgess and Marshall 2014). Even if natural environments are correlated, the evidence for stronger parental effects in more stable environments is mixed, and varies between traits (Walsh et al. 2016). Thus, there is a call for studies estimating parent and offspring fitness when the environmental predictability between generations are well known, and maternal effects are expected to be under selection (Burgess and Marshall 2014).

Direct experimental evidence for the evolution of positive anticipatory parental effects of an environmentally induced parental phenotype is currently lacking. On the other hand, if parents and offspring live in negatively correlated environments, the parental phenotype should not be inherited, but the environment is predictable and parents can still anticipate the offspring environment. As such, the evolution of a negative parental effect could be adaptive, a prediction that has recently received experimental support in a study by (Dey et al. 2016), who found that populations of the nematode *Caenorhabditis elegans* that evolved under strictly alternating hypoxia-normoxia conditions in every generation evolved a negative maternal provisioning effect. This suggests an adaptive benefit of maternal effects when the maternal environment is a strong cue for the offspring environment (in this case a perfect negative correlation), as predicted by theory (Uller et al. 2015; Proulx et al. 2017). More generally, theory also predicts that positive trans-generational correlations would result in the evolution of a positive parental effect and, importantly, if the environmental state is uncorrelated and unpredictable across generations, parental effects would be maladaptive and selected against (Lachmann and Jablonka 1996; Kuijper and Hoyle 2015; Leimar and McNamara 2015; Uller et al. 2015). The latter is considered a reason why adaptive parental effects are generally weak (Uller et al. 2013). However, these scenarios have not been investigated experimentally. Moreover, previous studies on evolution of parental effects under environmental heterogeneity have investigated only the case of non-overlapping generations (Dey et al. 2016). Most natural populations have however overlapping generations and age structure, which can influence evolution in both stable and heterogeneous environments, especially with respect to the evolution of life history strategies (Cotto and Ronce 2014; Ratikainen and Kokko 2019).

Taken together, environmental heterogeneity and environmental predictability over generations predict the adaptive value of parental effects (Lachmann and Jablonka 1996; Kuijper and Hoyle 2015; Uller et al. 2015). We set out to test this using experimental evolution in the dioecious free-living nematode *Caenorhabditis remanei*, adapting to different temperature regimes. Genetically heterogeneous populations, previously adapted to 20°C, were evolving for 30 generations in control conditions or adapting to 25°C, in either constant 25°C, increased warming to 25°C or a heterogeneous environment with fluctuating temperatures. We found positive anticipatory maternal effects on reproduction in populations evolving in environments that were predictable across generations. Moreover, we found the evolution of a reduced maternal effect on reproduction in heterogeneous environments where parent and offspring environments were not correlated during experimental evolution.

## Material and Methods

### Experimental evolution

As founder population, we used the wild-type SP8 strain of *C. remanei*, obtained from N. Timmermeyer at the Department of Biology, University of Tübingen, Germany. This strain was created by crossing three wild-type isolates of *C. remanei* (SB146, MY31, and PB206), harbour substantial standing genetic variation for life-history traits (Chen and Maklakov 2012; Lind et al. 2017), and has been lab-adapted to 20°C for 15 generations prior to setup of experimental evolution.

Experimental evolution was conducted for 30 generations in three climate cabinets; one set to 20°C, one to 25°C and one with a slowly increasing temperature (see below). Five experimental evolution regimes were used, and are summarized in Figure 1. *Control 20°C* was experiencing 20°C for 30 generations, and *Warm 25°C* was experiencing 25°C for 30 generations. *Increased warming* started in 20°C, the cabinet temperature was then raised by 0.1°C every 2.13 day (rounded to whole days) to reach 25°C the last day of experimental evolution. *Slow temperature cycles* spend their first 10 generations in 20°C, were then moved to the 25°C cabinet for 10 generations, to finish the last 10 generations in the 20°C cabinet. Finally, the *Fast temperature cycles* regime were moved between 20°C and 25°C every second generation, thus experiencing 14 temperature shifts.

**Figure 1.**
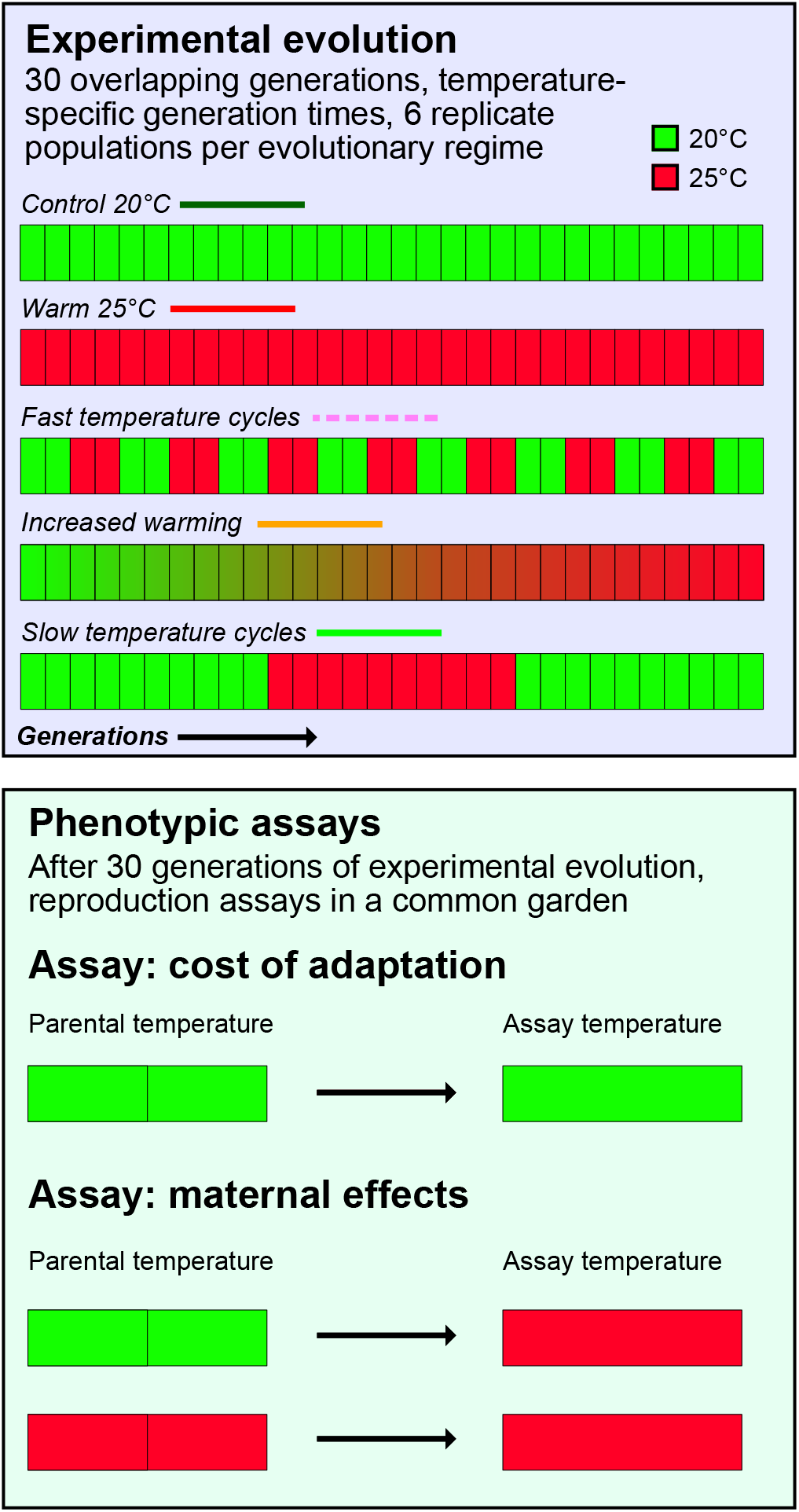
Overview of the experimental evolution regimes and the phenotypic assays. The five experimental evolution regimes (with 6 replicate populations) are outlined in the purple box. Green squares denotes 20°C and red squares 25°C. Each of the 30 small squares in each regime represents one generation. The phenotypic assays are presented in the green box, where parents are grown for two generations in either 20°C (green) or 25°C (red). Offspring are scored for age-specific fecundity, used to calculate total reproduction and daily growth factor λ.

Generation time in 20°C and 25°C was defined as the average difference in age between parents and offspring (Charlesworth 1994) and was calculated from the life-table of age-specific reproduction and survival following (McGraw and Caswell 1996) with trial data from the SP8 lines, and was 4.0 days in 20°C and 3.4 days in 25°C. This resulted 120 days of experimental evolution for *Control 20°C*, 114 days for *Slow temperature cycles*, 110 days for *Increased warming* and *Fast temperature cycles* and 101 days for *Warm 25°C*. For the two temperature cycle treatments, the worms spent shorter chronological time in 25°C than in 20°C, because of the faster generation time in 25°C. This ensured equal exposure to the two temperatures over biological time. Since we had no data for generation time in the intermediate temperatures between 20°C and 25°C, we did the simplifying assumption that the overall number of generations for the whole experiment would be similar in the *Increased warming* and *Fast temperature cycle* regimes. Therefore, the *Increased warming* regime was also run for 110 days, and the temperature increase was determined by the smallest temperature step the cabinet could be programmed (0.1°C).

Experimental evolution was conducted on 92 mm NGM-plates (Stiernagle 2006) and to combat infections the agar and bacterial LB also contained the antibiotics kanamycin and streptomycin, and the fungicide nystatin (Lionaki and Tavernarakis 2013; Lind et al. 2016). The plates were seeded with 2mL of the antibiotic resistant *E. coli* strain OP50-1 (pUC4K) obtained from J. Ewbank at the Centre d’Immunologie de Marseille-Luminy, France. To keep worm populations age-structured in overlapping generations, the populations were always kept in experimental growth face by cutting out a bit of agar containing 150 individuals of mixed ages and transferring this to freshly seeded plates. Transfer was conducted when needed (every 1-2 day), always before food was finished. Six independent replicate populations of each experimental evolution regime were set up, resulting in a total of 30 populations. All populations were expanded for two generations and frozen after 30 generations.

### Phenotypic assays

Before assays, worms were recover from freezing and grown 2 generations in common garden, each generation synchronized by bleaching, a standard procedure that kills all life-stages but eggs (Stiernagle 2006). The common garden temperature was 20°C or 25°C (see below).

Phenotypic assays were performed to test for local adaptation to the experimental evolution regime and the evolution of adaptive maternal effects, and are summarized in figure 1. We therefore carried out three assays, by varying parental temperature (the 2 generations of common garden after defrosting the populations) and testing temperature for offspring. The *20°C - 20°C assay* had both common garden and testing in 20°C. This is the environment the *Control 20°C* regime have experienced and tests for any cost of adaptation. Likewise, in the *25°C - 25°C assay* both parents and testing worms experience 25°C, which is the selective environment for *Warm 25°C* and very close to the final environment for *Increased warming*. Finally, the *20°C - 25°C assay* have 20°C as parental temperature, while the testing worms have their whole development in 25°C. This assay represents strong temperature fluctuations between generations, which is the selective environments for the *Fast temperature cycle* regime, and by comparing this assay to the *25°C - 25°C assay* we can estimate the importance of maternal effects on fitness when adapting to a novel environment.

The assays were initiated by synchronised egg-laying in the testing temperature by 40 females of each population. After 5h, females were killed by bleaching, and setup of L4 larvae was initiated 39h later (in 25°C) or 50h later (in 20°C), due to temperature-specific development time. The setup consisted of eight testing females per plate, together with the same number of background males from the SP8 line. Sex ratio was kept 1:1 throughout the experiment by adjusting the number of males to match the number of females present. Age-specific fecundity was measured by each day allowing the females 3h of egg-laying on an empty plate, where after the females were returned to a new plate (together with the males) and the number of hatched offspring on the egg-laying plate were killed with chloroform and counted two days later. The exact time the females were added to and removed from each plate was noted, and the number of offspring was corrected by exact number of minutes available for egg laying, and the number of females alive. Thus, we did not collect individual level data on total reproduction, but daily snapshot, in order to increase the number of individuals assayed and improve the reproduction estimate of each population. Daily reproduction was collected until reproduction had ceased. Four replicate plates of each population was set up, and for the *20°C - 20°C* and *20°C - 25°C* assays the replicates were evenly split between two climate cabinets per temperature, in order to separate cabinet and temperature effects. However, for logistical reasons, the *25°C - 25°C assay* was reduced. We excluded the *Slow temperature cycle* treatment from this assay, and unfortunately we lost two *Warm 25°C* populations during common garden (due to overcrowding and subsequent starving, which is known to induce epigenetic effects Rechavi et al. 2014) and therefore these populations were excluded), leaving us with four replicate population of this treatment. This resulted in 30 replicate populations and 960 female worms for the *20°C - 20°C* and *20°C - 25°C* assays, and 23 replicate populations and 736 female worms for the *25°C - 25°C assay*. A small number of plates were excluded from analyses, because of accidents during egg-laying or chloroforming: six plates were removed from the *20°C - 20°C* assay, four plates from the *20°C - 25°C* assay and six plates from the *25°C - 25°C* assay.

### Statistical analyses

The age-specific reproduction data was analysed as the total reproduction of each replicate plate, adjusted to the number of females present each day as well as rate-sensitive daily growth factor λ, which is equivalent to individual fitness (Brommer et al. 2002) but calculated per plate. λ encompasses the timing and number of offspring and is analogous to the intrinsic rate of population growth (Stearns 1992) and was calculated by constructing a population projection matrix for each plate, and then calculating the dominant eigenvalue from this matrix, following (McGraw and Caswell 1996). In the projection matrix, we set survival to 1 during the reproductive period, since any mortality during the early stages of the reproductive period (which is the most influential for λ) had non-natural causes (mainly worms climbing the wall of the plate where they dried out) and controlled for unequal number of worms by adjusting the reproductive value as described above. The first natural deaths in *C. remanei* are only observed long after the reproductive peak (Lind et al. 2016, 2017). Since population size and age-structure was kept constant during the experimental evolution, rate sensitive fitness is maximised during experimental evolution and λ is therefore the most appropriate fitness measure for this study (Mylius and Diekmann 1995).

All statistical analyses were done using *R 3.6.1*, and models were implemented using the *lme4* package (Bates et al. 2015). Significance tests were performed using the *lmerTest* package (Kuznetsova et al. 2017), and contrasts were analysed using the *emmeans* package. Total reproduction and daily growth factor λ in 20°C was analysed in separate mixed effect models with experimental evolution regime as fixed effect and population and cabinet as random effects. Both response variables were log-transformed before analysis. In 25°C, λ and total reproduction was analysed with experimental evolution regime and parental temperature as crossed fixed effects, and replicate population as random effect. Models were constructed with a random slope (parental temperature) in addition to random intercept. Since the *25°C - 25°C assay* was conducted in only one cabinet, and moreover the *Slow temperature cycle* treatment was not run, the random effect of cabinet was excluded from the parental temperature models, as was the *Slow temperature cycle* treatment.

## Results

The predictions and results for maternal effects are summarized in Table 1. For fitness (daily growth factor λ) in 25°C, we found a significant experimental evolution regime × parental temperature interaction (Regime: *F*_3, 20.2_ = 0.493, p = 0.691; Parental temperature: *F*_1, 18.9_ = 0.135, p = 0.718, Regime × Parental temperature: *F*_3, 18.7_ = 4.249, p = 0.019). This interaction was caused by significantly opposite slope for *Fast temperature cycles* compared to *Increased warming*, with highest fitness in 25°C for *Fast temperature cycles* when parents were grown in 20°C, while highest fitness in 25°C for *Increased warming* was achieved when their parents were also grown in 25°C. For *Control 20°C* and *Warm 25°C*, fitness in 25°C did not significantly differ between the parental temperatures (Figure 2A, suppl. table 1-2).

**Table 1.** Summary of the predictions and findings of the evolution of maternal effects based upon the stability and predictability between parent and offspring environment in the five experimental evolution regimes.

| Experimental evolution regime | Environment predictability over generations | Prediction | Finding: reproduction | Finding: daily growth factor $\lambda$ |
| --- | --- | --- | --- | --- |
| *Control 20°C* | Stable, predictable | Either positive or negative maternal effect (inherit maternal phenotype or reduce phenotypic variance) | Positive maternal effect | No maternal effect |
| *Warm 25°C* | Stable, predictable | Either positive or negative maternal effect (inherit maternal phenotype or reduce phenotypic variance) | Positive maternal effect | No maternal effect |
| *Increased warming* | Slowly changing, predictable | Positive maternal effect (inherit maternal phenotype) | Positive maternal effect | Positive maternal effect |
| *Slow temp. cycles* | Slowly changing, predictable | Positive maternal effect (inherit maternal phenotype) | Not tested | Not tested |
| *Fast temp. cycles* | Unpredictable | No maternal effect | No maternal effect | Negative maternal effect |

**Figure 2.**
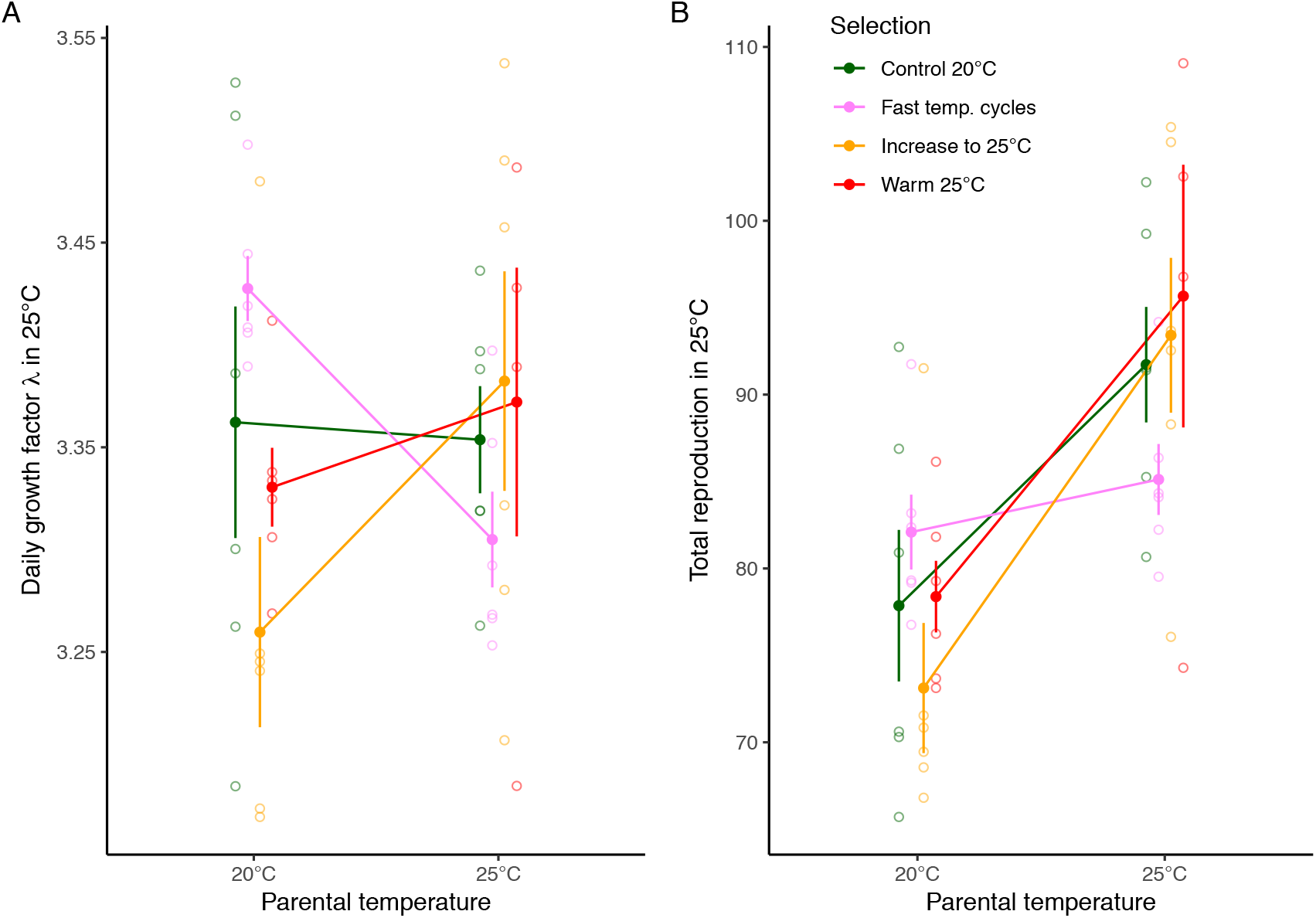
The effect of parental temperature on fitness and reproduction in 25°C. Daily growth factor λ (A) and total reproduction (B) in 25°C when parent were raised for 2 generations in either 20°C or 25°C. Symbols represent experimental evolution regime (mean ± SE calculated from population means). Open symbols represent the mean of each replicate population. *Control 20°C* and *Warm 25°C* have spent 30 generations in 20°C or 25°C, respectively. *Increased warming* has been subjected to slowly increased temperatures, starting in 20°C and reaching 25°C at generation 30. *Fast temperature* cycles have spent two generations in 20°C, followed by two generations in 25°C, this cycle has then been repeated for 30 generations.

For total reproduction in 25°C, the experimental evolution regime × parental temperature interaction was just outside the significance threshold (Regime: *F*_3, 19.7_ = 0.385, p = 0.765; Parental temperature: *F*_1, 18.8_ = 35.659, p < 0.001, Regime × Parental temperature: *F*_3, 18.6_ = 3.071, p = 0.053). Nevertheless, the planned post-hoc comparisons showed that there was no effect of parental exposure to 25°C for *Fast temperature cycles*, while all other regimes had higher total reproduction in 25°C if parents were also exposed to 25°C (Figure 2B, suppl. table 3-4).

Finally, we found a fitness cost of adaptation in many of the regimes, since *Fast temperature cycles* as well *Slow temperature cycles* had significantly reduced daily growth factor (λ) in 20°C relative to the *Control 20°C* regime (Regime: *F*_4, 24.6_ = 4.955, p = 0.005, Figure 3B, suppl. table 5-6). For the *Increased warming* and *Warm 25°C* regimes, the effect was just outside significance. The cost was however not detected in total reproduction (*F*_4, 24.8_ = 0.574, p = 0.684, Figure 3A, suppl. table 7-8).

**Figure 3.**
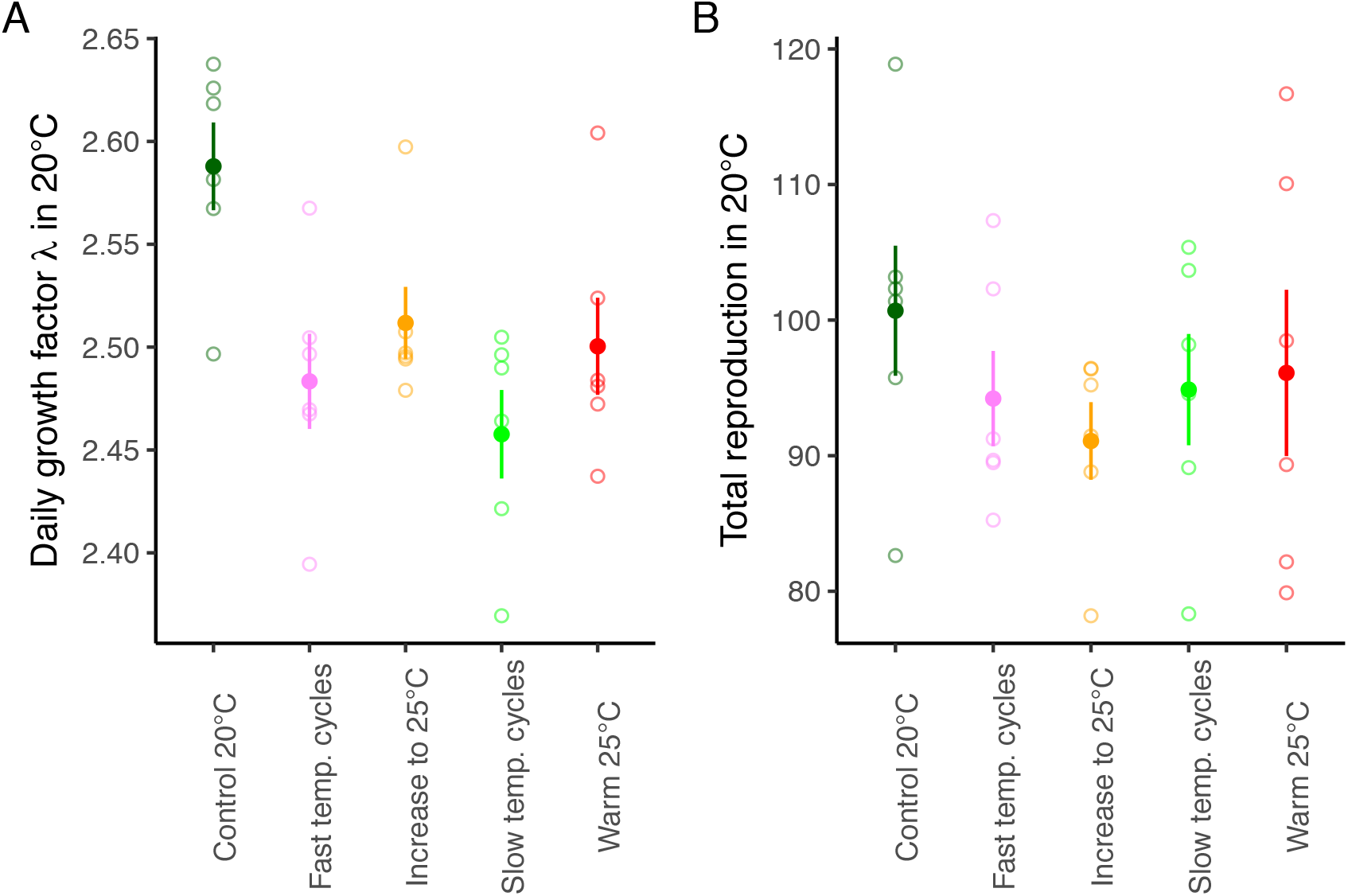
Reproduction in original environment. Daily growth factor λ (A) and total reproduction (B) in 20°C when parents are also grown for two generations in 20°C. Symbols represent experimental evolution regime (mean ± SE calculated from population means). Open symbols represent the mean of each replicate population.

## Discussion

The degree of environmental variation can influence the expression of phenotypes (Moran 1992; Uller et al. 2015; Proulx et al. 2017). In addition to genetic specialisation, the phenotype can match the environment either by phenotypic plasticity, where the offspring matches its development to the current environment, or by anticipatory parental effects, where the offspring non-genetically inherit the parents phenotype to match its environment, thus improving offspring performance if parent and offspring environments are matching (Marshall and Uller 2007; Donelson et al. 2017). While within-generation phenotypic plasticity is common (DeWitt and Scheiner 2004), parental effects seem, in contrast, to be generally weak (Uller et al. 2013), despite some well known examples (Gustafsson et al. 2005; Galloway and Etterson 2007; Jensen et al. 2014; Kishimoto et al. 2017; Ryu et al. 2018; Toyota et al. 2019). Since trans-generational effects should evolve only if parents can accurately predict offspring environments (Kuijper and Hoyle 2015; Leimar and McNamara 2015; Uller et al. 2015; Dury and Wade 2020), it is possible that environments generally are not highly correlated between generations, thus explaining why such anticipatory effects are uncommon (Uller et al. 2013). However, the environmental predictability is seldom investigated in studies of parental effects (Burgess and Marshall 2014). Therefore, we investigated whether the degree of temporal environmental variation, as well as the predictability between parent and offspring environment, influenced the evolution of maternal effects.

We found that the presence of environmental variation mediated the evolution of maternal effects on reproduction and daily growth factor λ in *C. remanei* nematode worms adapting to a novel and stressful warm temperature (25°C) for 30 generations (see predictions and findings summarized in Table 1). For all populations evolving in stable or slowly increasing temperature (*Control 20°C*, *Warm 25°C*, *Increased warming*), a strong positive maternal effect on reproductive output resulted in an increased offspring production in 25°C when the parents were also cultured in 25°C and not in 20°C. Since these populations have evolved in environments that are relatively predictable over generations, trans-generational effects are adaptive and predicted by theory (Mousseau and Fox 1998; Kuijper and Hoyle 2015; Leimar and McNamara 2015; Uller et al. 2015). This result is also in agreement with a recent study by Dey et al. (2016) who found the evolution of an anticipatory negative maternal effect in *C. elegans* evolving in perfectly negatively correlated (and thus predictable) environments. Our finding of positive anticipatory maternal effects in positively correlated environments highlights the importance of experimental evolution studies with known environmental predictability to study the evolution of adaptive trans-generational effects.

In contrast to the predictable environments, the *Fast temperature cycles* populations evolved in a fluctuating environment where the temperature changed every second generation. Thus, the next generation would with equal likelihood be exposed to the same or a different temperature as the parents, resulting in zero correlation between parent and offspring environments. In this unpredictable environment, trans-generational effects are not considered adaptive (Mousseau and Fox 1998; Uller et al. 2015), and, in agreement with the theory, we found a loss of the positive maternal effect, since the reproductive output in 25°C of this regime was independent of the environment their parents. It should be noted that while the interaction between experimental evolution regime and parental temperature was non-significant (p = 0.053) for total reproduction, the planned comparisons showed clearly that maternal effects were strong and positive in all regimes except *Fast temperature cycles*, where they were far from significant (supplementary table 4). The adaptive value of these differences in parental effects is illustrated in the daily growth factor λ of the different experimental evolution regimes. *Fast temperature cycles* populations had highest λ in 25°C only when the parents were cultured for two generations in 20°C, a situation mimicking the fluctuating environments they were exposed to during evolution. Although adaptive, this should be defined as negative maternal effect on λ, which is not predicted by theory. Thus, maternal effects on total reproduction and λ does not follow the same pattern for the *Fast temperature cycles*, but importantly, none of the measures show the positive maternal effects present in regimes from more predictable environments (figure 2). In contrast, populations adapting to slowly changing warm temperatures (*Increased warming*) improved λ when parents were also cultured in 25°C, and thus showed a positive maternal effect on λ. Interestingly, the two stable environments (*Control 20°C*, *Warm 25°C*), who showed a positive maternal effect on reproduction did not show any maternal effect for λ (although there was a strong positive trend for *Warm 25°C*). Since stable environments are predicted to not show a positive maternal effect when fully adapted (Kuijper and Hoyle 2015), the loss of maternal effect for λ but not for total reproduction may be a result of ongoing genetic specialisation, replacing transient maternal effects. Whether the negative maternal effect on fitness in the *Fast temperature cycles* is a result of the fact that the assays with parents and offspring in 25°C represent a stability not experienced evolutionary by the *Fast temperature cycles* regime (two generations common garden for the parents and one generation offspring testing equals to three generations in the same environment), or is a general response when parent and offspring environment is matching is unknown. The discrepancy between total reproduction and daily growth factor λ is illustrated by different patterns of age-specific reproduction in the two temperatures (Figure S1). While reproduction overall is high in 25°C - 25°C, with peak reproduction spanning the first two days of adulthood, the reproductive peak is higher and earlier in the 20°C - 25°C, but reproduction also declines faster. The fast cycle regime perform well during the first day of adulthood in the 20°C - 25°C assay, where after its age-specific reproduction quickly drops, which affect total reproduction much more than rate-sensitive fitness measures such as λ, that weight early reproduction higher than late reproduction (Brommer et al. 2002).

In addition, for the two cyclic regimes we also found a fitness cost of adaptation, since they had lower λ in the original environment (20°C), compared to the *Control 20°C* regime. The same trend of a fitness cost was shown also for the *Increased warming* and *Warm 25°C* regimes, but was just outside significance. A cost of adaptation was however not significant for total reproduction, highlighting the fitness importance of the increased early reproduction during the two first days of adulthood in the *Control 20°C* populations (Figure S2). A cost of adaptation, manifested as antagonistic environmental pleiotropy, is previously found in studies of antibiotic resistance, where the resistance comes with a cost in the original environment (Gifford et al. 2018). Similar costs have also been found in experimental evolution studies (Kassen 2002), also when adapting to different temperature regimes in insects (Tobler et al. 2015; Berger et al. 2018). Moreover, although *Control 20°C* showed a positive maternal effect on reproduction in 25°C, maternal exposure to 25°C did not improve their individual fitness, suggesting that an evolutionary history in 25°C is needed for maximum fitness in this temperature.

While the positive maternal effect present in both stable (*Control 20°C*, *Warm 25°C*) and slowly increasing (*Increased warming*) temperature regimes is anticipated when parent can predict offspring environments (Lachmann and Jablonka 1996; Uller et al. 2015), stable environments are actually predicted to select for negative trans-generational effects (Hoyle and Ezard 2012; Kuijper and Hoyle 2015). When a population is well adapted to a stable optimum, a negative maternal effect reduces phenotypic variance between generations. However, we find no support for this prediction, since none of the treatments in stable environments (*Control 20°C*, *Warm 25°C*) showed a negative maternal effect, even after 30 generations in stable conditions. It is however possible that these regimes still show transient dynamics, since a positive trans-generational effect is predicted to evolve as a transient response when experiencing a novel environment (Kuijper and Hoyle 2015), in a similar way to how phenotypic plasticity is predicted to aid adaptation to new environments (Price et al. 2003; Lande 2009; Chevin et al. 2010; Coulson et al. 2017). Moreover, while these lines show a positive maternal effect for total reproduction, they show no significant maternal effect for λ, which could indicate an ongoing loss of maternal effects in stable environments, in line with theory. It could possibly be argued that the positive trans-generational effect is non-adaptive, caused by the worms being maladapted in 20°C and therefore producing low-quality offspring in this temperature. However, the fact that the *Control 20°C* regime, who have highest fitness in 20°C and who have not experienced 25°C for at least 45 generations show positive maternal effects on reproduction of parental exposure to 25°C argues against a non-adaptive explanation and instead reinforces the view that all regimes from stable and slowly changing environment has an adaptive positive maternal effect. In agreement with this, exposing a laboratory strain of *C. elegans* to 25°C (Schott et al. 2014) or to starvation (Rechavi et al. 2014) results in heritable changes in gene expression, directed by inherited small RNA (Rechavi et al. 2014; Schott et al. 2014; Houri-Zeevi and Rechavi 2017), which further suggests that transgenerational plasticity such as maternal effects and epigenetic inheritance is important in *Caenorhabditis* nematodes.

We found that the environmental predictability between generations is driving the evolution of anticipatory maternal effects. While stable and slowly changing environments select for positive anticipatory maternal effects, environments that fluctuate with no correlation between generations select against parental influence on offspring phenotype. This is the first empirical study that investigates the evolutionary loss of anticipatory maternal effect, which follow theoretical predictions (Kuijper and Hoyle 2015; Uller et al. 2015) and suggest that one reason for the weak effects of parent environment on offspring phenotype in natural systems (Uller et al. 2013) could be that natural environments are not always predictable across generations. While most examples of positive parental effects comes from systems with short lifecycles such as nematodes (Kishimoto et al. 2017), daphniids (Gustafsson et al. 2005; Toyota et al. 2019), annelids (Jensen et al. 2014) and herbs (Galloway and Etterson 2007), it is also been found in fish where the generation time span years (Ryu et al. 2018). However, when investigating maternal effects in *Daphnia* from natural populations with different degree of variation in predation intensity, (Walsh et al. 2016) found some support for stronger positive maternal effects in population from more stable environments, but the effect differed between traits and no effect was found on reproduction. Nevertheless, these types of studies where the environmental predictability is known are vital for our understanding of the selection pressures resulting in the presence or absence of adaptive maternal effects in natural populations (Burgess and Marshall 2014), where deconstruction of environmental predictability to seasonality and environmental colour noise (following Marshall and Burgess 2015) is a very promising approach.

On-going climate change is not only resulting in warmer temperatures, but also increased temperature variation, which can impact both the ecology and evolution of populations and species (Vasseur et al. 2014; Vázquez et al. 2017), and trans-generational acclimation can be an important response to deal with a warming climate (Donelson et al. 2017; Ryu et al. 2018). We show that environmental heterogeneity drives the evolution of maternal effects, and support the theoretical predictions (Lachmann and Jablonka 1996; Uller et al. 2015) that the predictability between parent and offspring environment is the driver of the evolution of transgenerational plasticity.

## Author contributions

MIL and AAM designed the experiment, MIL, JA, HC, TK and TL performed experimental evolution, MIL, MKZ and JA performed phenotypic assays, MIL analysed the data. MIL drafted the manuscript together with AAM. All authors contributed to the revision of the manuscript.

## Acknowledgement

This work was funded by the Swedish Research Council [grant number 2016-05195 to MIL, grant number 621-2013-4828 to AAM] and the European Research Council [grants AGINGSEXDIFF and GERMLINEAGEINGSOMA to AAM]. We thank Frank Johansson, Anssi Laurila, Irja I. Ratikainen and Claus Rüffler for discussions.

## Data sharing

If accepted, data will be deposited at *Dryad*.

## Literature cited

Bates, D., M. Mächler, B. M. Bolker, and S. C. Walker. 2015. Fitting linear mixed-effects models using lme4. J. Stat. Softw. 67:1–48.

Berger, D., J. Stångberg, J. Baur, and R. J. Walters. 2018. A universal temperature-dependence of mutational fitness effects. bioRxiv 268011.

Bonduriansky, R., and T. Day. 2009. Nongenetic inheritance and its evolutionary implications. Annu. Rev. Ecol. Evol. Syst. 40:103–125.

Brommer, J. E., J. Merilä, and H. Kokko. 2002. Reproductive timing and individual fitness. Ecol. Lett. 5:802–810.

Burgess, S. C., and D. J. Marshall. 2014. Adaptive parental effects: the importance of estimating environmental predictability and offspring fitness appropriately. Oikos 123:769–776.

Charlesworth, B. 1994. Evolution in age-structured populations. 2nd ed. Cambridge University Press, New York, NY, USA.

Chen, H., and A. A. Maklakov. 2012. Longer life span evolves under high rates of condition-dependent mortality. Curr. Biol. 22:2140–2143.

Chevin, L.-M., R. Lande, and G. M. Mace. 2010. Adaptation, plasticity, and extinction in a changing environment: towards a predictive theory. PLoS Biol 8:e1000357.

Cotto, O., and O. Ronce. 2014. Maladaptation as a source of senescence in habitats variable in space and time. Evolution 68:2481–2493.

Coulson, T., B. E. Kendall, J. Barthold, F. Plard, S. Schindler, A. Ozgul, and J.-M. Gaillard. 2017. Modeling adaptive and nonadaptive responses of populations to environmental change. Am. Nat. 190:313–336.

DeWitt, T. J., and S. M. Scheiner. 2004. Phenotypic plasticity: functional and conceptual approaches. Oxford University Press, New York, NY, USA.

Dey, S., S. R. Proulx, and H. Teotónio. 2016. Adaptation to temporally fluctuating environments by the evolution of maternal effects. PLoS Biol. 14:e1002388.

Donelson, J. M., S. Salinas, P. L. Munday, and L. N. S. Shama. 2017. Transgenerational plasticity and climate change experiments: Where do we go from here? Glob. Change Biol. 24:13–34.

Dury, G. J., and M. J. Wade. 2020. When mother knows best: A population genetic model of transgenerational versus intragenerational plasticity. J. Evol. Biol. 33:127–137.

Galloway, L. F., and J. R. Etterson. 2007. Transgenerational plasticity is adaptive in the wild. Science 318:1134–1136.

Gifford, D. R., R. Krašovec, E. Aston, R. V. Belavkin, A. Channon, and C. G. Knight. 2018. Environmental pleiotropy and demographic history direct adaptation under antibiotic selection. Heredity 121:438–448.

Gustafsson, S., K. Rengefors, and L.-A. Hansson. 2005. Increased consumer fitness following transfer of toxin tolerance to offspring via maternal effects. Ecology 86:2561–2567.

Hollander, J., E. Snell-Rood, and S. Foster. 2015. New frontiers in phenotypic plasticity and evolution. Heredity 115:273–275.

Houri-Zeevi, L., and O. Rechavi. 2017. A matter of time: small RNAs regulate the duration of epigenetic inheritance. Trends Genet. 33:46–57.

Hoyle, R. B., and T. H. G. Ezard. 2012. The benefits of maternal effects in novel and in stable environments. J. R. Soc. Interface 9:2403–2413.

Jensen, N., R. M. Allen, and D. J. Marshall. 2014. Adaptive maternal and paternal effects: gamete plasticity in response to parental stress. Funct. Ecol. 28:724–733.

Kassen, R. 2002. The experimental evolution of specialists, generalists, and the maintenance of diversity: Experimental evolution in variable environments. J. Evol. Biol. 15:173–190.

Kishimoto, S., M. Uno, E. Okabe, M. Nono, and E. Nishida. 2017. Environmental stresses induce transgenerationally inheritable survival advantages via germline-to-soma communication in *Caenorhabditis elegans*. Nat. Commun. 8:14031.

Kuijper, B., and R. B. Hoyle. 2015. When to rely on maternal effects and when on phenotypic plasticity? Evolution 69:950–968.

Kuznetsova, A., P. B. Brockhoff, and R. H. B. Christensen. 2017. lmerTest package: tests in linear mixed effects models. J. Stat. Softw. 82:1–26.

Lachmann, M., and E. Jablonka. 1996. The inheritance of phenotypes: an adaptation to fluctuating environments. J. Theor. Biol. 181:1–9.

Lande, R. 2009. Adaptation to an extraordinary environment by evolution of phenotypic plasticity and genetic assimilation. J. Evol. Biol. 22:1435–1446.

Leimar, O., and J. M. McNamara. 2015. The evolution of transgenerational integration of information in heterogeneous environments. Am. Nat. 185:E55–E69.

Lind, M. I., H. Chen, S. Meurling, A. C. Guevara Gil, H. Carlsson, M. K. Zwoinska, J. Andersson, T. Larva, and A. A. Maklakov. 2017. Slow development as an evolutionary cost of long life. Funct. Ecol. 31:1252–1261.

Lind, M. I., and F. Spagopoulou. 2018. Evolutionary consequences of epigenetic inheritance. Heredity 121:205–209.

Lind, M. I., M. K. Zwoinska, S. Meurling, H. Carlsson, and A. A. Maklakov. 2016. Sex-specific trade-offs with growth and fitness following lifespan extension by rapamycin in an outcrossing nematode, *Caenorhabditis remanei*. J. Gerontol. A. Biol. Sci. Med. Sci. 71:882–890.

Lionaki, E., and N. Tavernarakis. 2013. Assessing aging and senescent decline in *Caenorhabditis elegans*: cohort survival analysis. Pp. 473–484 *in* L. Galluzzi, I. Vitale, O. Kepp, and G. Kroemer, eds. Cell Senescence. Humana Press, New York.

Marshall, D. J., and S. C. Burgess. 2015. Deconstructing environmental predictability: seasonality, environmental colour and the biogeography of marine life histories. Ecol. Lett. 18:174–181.

Marshall, D. J., and T. Uller. 2007. When is a maternal effect adaptive. Oikos 116:1957–1963.

McGraw, J. B., and H. Caswell. 1996. Estimation of individual fitness from life-history data. Am. Nat. 147:47–64.

Moran, N. A. 1992. The evolutionary maintenance of alternative phenotypes. Am. Nat. 139:971–989.

Mousseau, T. A., and C. W. Fox. 1998. The adaptive significance of maternal effects. Trends Ecol. Evol. 13:403–407.

Mylius, S. D., and O. Diekmann. 1995. On evolutionarily stable life histories, optimization and the need to be specific about density dependence. Oikos 74:218–224.

Perez, M. F., and B. Lehner. 2019. Intergenerational and transgenerational epigenetic inheritance in animals. Nat. Cell Biol. 21:143.

Price, T. D., A. Qvarnström, and D. E. Irwin. 2003. The role of phenotypic plasticity in driving genetic evolution. Proc. R. Soc. Lond. Ser. B-Biol. Sci. 270:1433–1440.

Proulx, S. R., H. Teotónio, R. Bonduriansky, and Y. Michalakis. 2017. What kind of maternal effects can be selected for in fluctuating environments? Am. Nat. 189:E118–E137.

Ratikainen, I. I., and H. Kokko. 2019. The coevolution of lifespan and reversible plasticity. Nat. Commun. 10:538.

Rechavi, O., L. Houri-Ze’evi, S. Anava, W. S. S. Goh, S. Y. Kerk, G. J. Hannon, and O. Hobert. 2014. Starvation-induced transgenerational inheritance of small RNAs in *C. elegans*. Cell 158:277–287.

Rivoire, O., and S. Leibler. 2014. A model for the generation and transmission of variations in evolution. Proc. Natl. Acad. Sci. 111:E1940–E1949.

Ryu, T., H. D. Veilleux, J. M. Donelson, P. L. Munday, and T. Ravasi. 2018. The epigenetic landscape of transgenerational acclimation to ocean warming. Nat. Clim. Change 8:504–509.

Schott, D., I. Yanai, and C. P. Hunter. 2014. Natural RNA interference directs a heritable response to the environment. Sci. Rep. 4:7387.

Simons, A. M. 2011. Modes of response to environmental change and the elusive empirical evidence for bet hedging. Proc. R. Soc. Lond. B Biol. Sci. 278:1601–1609.

Stearns, S. C. 1992. The evolution of life histories. Oxford University Press, New York, NY, USA.

Stiernagle, T. 2006. Maintenance of *C. elegans*. WormBook Online Rev. C Elegans Biol., doi: 10.1895/wormbook.1.101.1.

Tobler, R., J. Hermisson, and C. Schlötterer. 2015. Parallel trait adaptation across opposing thermal environments in experimental *Drosophila melanogaster* populations. Evolution 69:1745–1759.

Toyota, K., M. C. Cuenca, V. Dhandapani, A. Suppa, V. Rossi, J. K. Colbourne, and L. Orsini. 2019. Transgenerational response to early spring warming in *Daphnia*. Sci. Rep. 9:4449.

Uller, T., S. English, and I. Pen. 2015. When is incomplete epigenetic resetting in germ cells favoured by natural selection? Proc R Soc B 282:20150682.

Uller, T., S. Nakagawa, and S. English. 2013. Weak evidence for anticipatory parental effects in plants and animals. J. Evol. Biol. 26:2161–2170.

Vasseur, D. A., J. P. DeLong, B. Gilbert, H. S. Greig, C. D. G. Harley, K. S. McCann, V. Savage, T. D. Tunney, and M. I. O’Connor. 2014. Increased temperature variation poses a greater risk to species than climate warming. Proc. R. Soc. B Biol. Sci. 281:20132612.

Vázquez, D. P., E. Gianoli, W. F. Morris, and F. Bozinovic. 2017. Ecological and evolutionary impacts of changing climatic variability. Biol. Rev. 92:22–42.

Walsh, M. R., T. Castoe, J. Holmes, M. Packer, K. Biles, M. Walsh, S. B. Munch, and D. M. Post. 2016. Local adaptation in transgenerational responses to predators. Proc R Soc B 283:20152271.

