## Supplementary materials for "Environmental variation mediates the evolution of anticipatory parental effects"

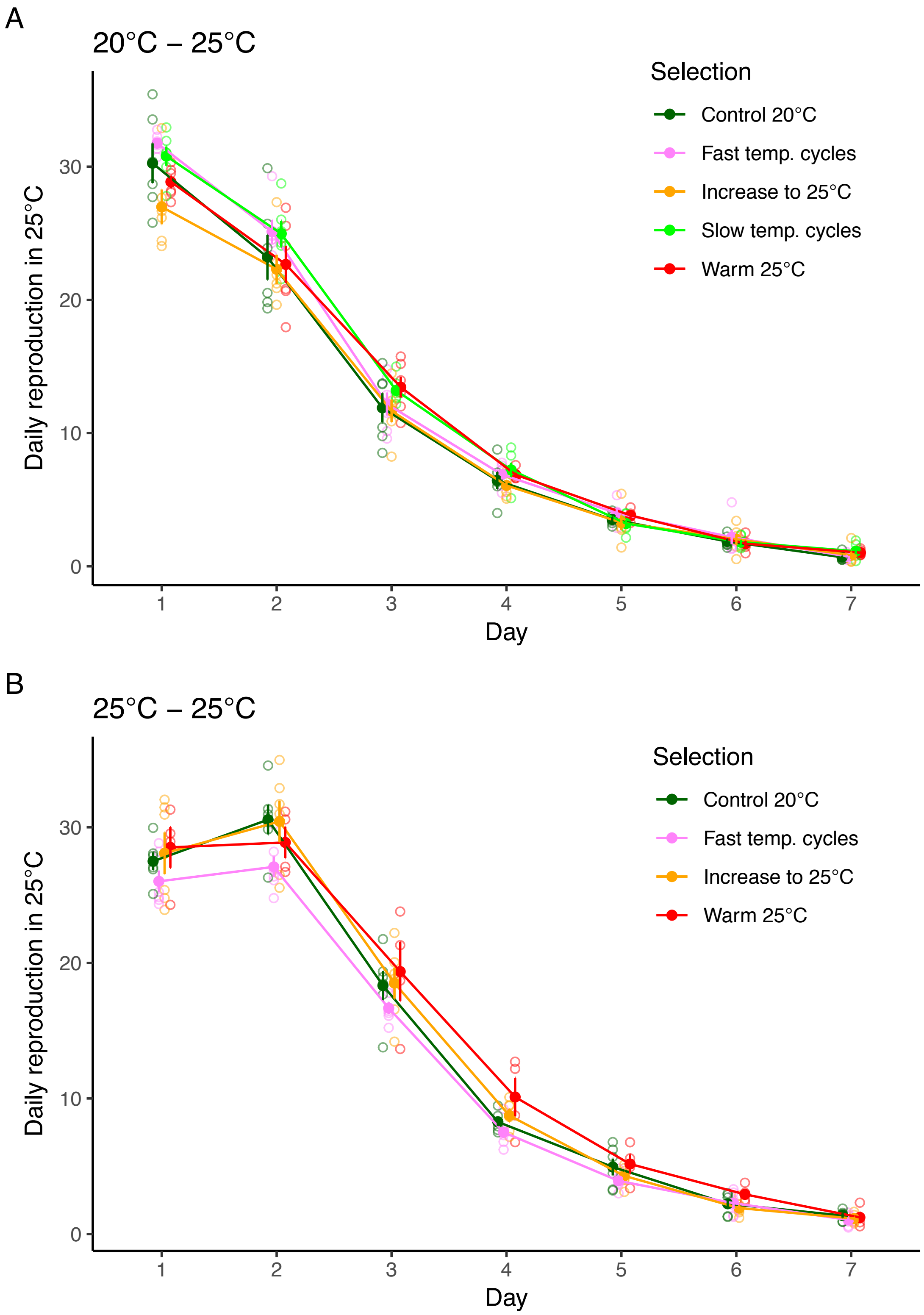


**Supplementary figure 1.** Daily reproduction in 25°C when parents are (A) grown for two generations in 20°C or (B) grown for two generations in 25°C. Symbols represent experimental evolution regime (mean ± SE calculated from line means). Open symbols represent the mean of each replicate line.

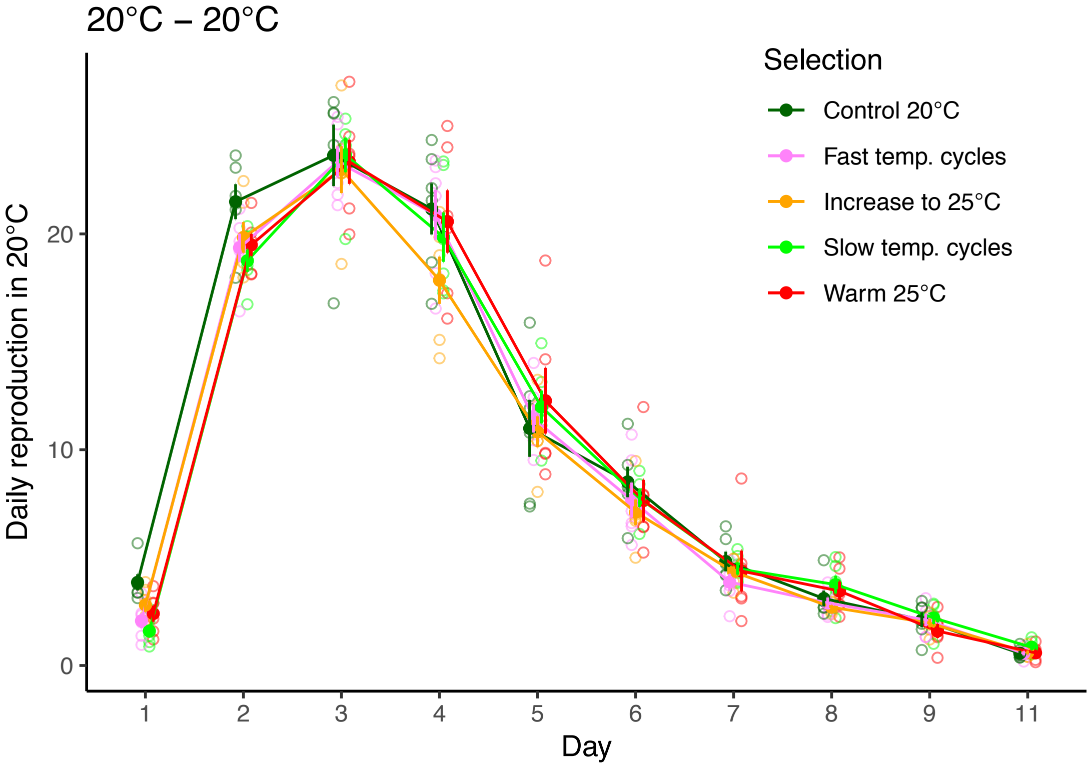


**Supplementary figure 2.** Daily reproduction in 20°C when parents are also grown for two generations in 20°C. Symbols represent experimental evolution regime (mean ± SE calculated from line means). Open symbols represent the mean of each replicate line.

**Supplementary table 1.** Daily growth factor λ in 25°C. Estimated experimental evolution regime means, with associated 95% confidence limits.

| Regime | Parental temp. | emmean | SE | df | Lower CL | Upper CL |
|---|---|---|---|---|---|---|
| Fast temp. cycles | 20°C | 1.23 | 0.0124 | 21.8 | 1.21 | 1.26 |
| Fast temp. cycles | 25°C | 1.19 | 0.0119 | 17 | 1.17 | 1.22 |
| Control 20°C | 20°C | 1.21 | 0.0118 | 18.9 | 1.19 | 1.24 |
| Control 20°C | 25°C | 1.21 | 0.0124 | 18.6 | 1.18 | 1.23 |
| Incr. warming | 20°C | 1.18 | 0.012 | 19.9 | 1.16 | 1.21 |
| Incr. warming | 25°C | 1.22 | 0.0119 | 17 | 1.19 | 1.24 |
| Warm 25°C | 20°C | 1.2 | 0.0118 | 18.9 | 1.18 | 1.23 |
| Warm 25°C | 25°C | 1.22 | 0.0152 | 19.4 | 1.19 | 1.25 |

**Supplementary table 2.** Daily growth factor λ in 25°C. Post-hoc contrasts over temperatures, investigating the significant Regime × Parental temperature interaction from the full model.

| Regime | Contrast | Estimate | SE | df | t-ratio | p-value |
|---|---|---|---|---|---|---|
| Fast temp. cycles | 20°C - 25°C | 0.037 | 0.0151 | 18.8 | 2.481 | 0.023 |
| Control 20°C | 20°C - 25°C | 0.002 | 0.0149 | 18.1 | 0.138 | 0.892 |
| Incr. warming | 20°C - 25°C | -0.036 | 0.0147 | 17.4 | -2.416 | 0.027 |
| Warm 25°C | 20°C - 25°C | -0.015 | 0.0173 | 21.1 | -0.879 | 0.389 |

**Supplementary table 3.** Total reproduction in 25°C. Estimated experimental evolution regime means, with associated 95% confidence limits.

| Regime | Parental temp. | emmean | SE | df | Lower CL | Upper CL |
|---|---|---|---|---|---|---|
| Fast temp. cycles | 20°C | 4.4 | 0.045 | 21.8 | 4.31 | 4.5 |
| Fast temp. cycles | 25°C | 4.43 | 0.044 | 17 | 4.34 | 4.53 |
| Control 20°C | 20°C | 4.34 | 0.042 | 18.9 | 4.25 | 4.43 |
| Control 20°C | 25°C | 4.51 | 0.045 | 18.6 | 4.42 | 4.61 |
| Incr. warming | 20°C | 4.27 | 0.043 | 19.9 | 4.18 | 4.36 |
| Incr. warming | 25°C | 4.52 | 0.044 | 17 | 4.42 | 4.61 |
| Warm 25°C | 20°C | 4.35 | 0.042 | 18.9 | 4.26 | 4.44 |
| Warm 25°C | 25°C | 4.55 | 0.055 | 19.6 | 4.43 | 4.67 |

**Supplementary table 4.** Total reproduction in 25°C. Post-hoc contrasts over temperatures, investigating the significant Regime × Parental temperature interaction from the full model.

| Regime | Contrast | Estimate | SE | df | t-ratio | p-value |
|---|---|---|---|---|---|---|
| Fast temp. cycles | 20°C - 25°C | -0.030 | 0.0529 | 18.7 | -0.56 | 0.582 |
| Control 20°C | 20°C - 25°C | -0.177 | 0.0524 | 18 | -3.378 | 0.003 |
| Incr. warming | 20°C - 25°C | -0.243 | 0.0516 | 17.3 | -4.699 | <0.001 |
| Warm 25°C | 20°C - 25°C | -0.200 | 0.0613 | 20.8 | -3.263 | 0.004 |

**Supplementary table 5.** Daily growth factor λ in 20°C. Estimated experimental evolution regime means, with associated 95% confidence limits.

| Selection | emmean | SE | df | Lower CL | Upper CL |
|---|---|---|---|---|---|
| Control 20°C | 0.95 | 0.00963 | 22.4 | 0.93 | 0.97 |
| Fast temp. cycles | 0.909 | 0.00923 | 19.1 | 0.89 | 0.928 |
| Incr. warming | 0.92 | 0.00943 | 20.6 | 0.901 | 0.94 |
| Slow temp. cycles | 0.899 | 0.00923 | 19.1 | 0.879 | 0.918 |
| Warm 25°C | 0.916 | 0.00932 | 19.8 | 0.897 | 0.936 |

**Supplementary table 6.** Daily growth factor λ in 20°C. Post-hoc contrasts.

| Contrast | Estimate | SE | df | t-ratio | p-value |
|---|---|---|---|---|---|
| Control - Fast | 0.04118 | 0.0122 | 25.8 | 3.383 | 0.0178 |
| Control - Incr. | 0.02982 | 0.0123 | 26.9 | 2.421 | 0.1403 |
| Control - Slow | 0.05137 | 0.0122 | 25.8 | 4.22 | 0.0023 |
| Control - Warm | 0.03386 | 0.0122 | 26.3 | 2.764 | 0.0708 |
| Fast - Incr. | -0.01136 | 0.012 | 24.5 | -0.946 | 0.8761 |
| Fast - Slow | 0.01019 | 0.0119 | 23.4 | 0.86 | 0.9086 |
| Fast - Warm | -0.00733 | 0.0119 | 23.9 | -0.614 | 0.9714 |
| Incr. - Slow | 0.02155 | 0.012 | 24.5 | 1.794 | 0.3995 |
| Incr. - Warm | 0.00404 | 0.0121 | 25 | 0.334 | 0.9971 |
| Slow - Warm | -0.01752 | 0.0119 | 23.9 | -1.468 | 0.5917 |

**Supplementary table 7.** Total reproduction in 20°C. Estimated experimental evolution regime means, with associated 95% confidence limits.

| Selection | emmean | SE | df | Lower CL | Upper CL |
|---|---|---|---|---|---|
| Control 20°C | 4.6 | 0.0477 | 26.4 | 4.5 | 4.7 |
| Fast temp. cycles | 4.54 | 0.0462 | 23.4 | 4.44 | 4.63 |
| Incr. warming | 4.5 | 0.047 | 24.7 | 4.41 | 4.6 |
| Slow temp. cycles | 4.54 | 0.0462 | 23.4 | 4.45 | 4.64 |
| Warm 25°C | 4.55 | 0.0466 | 24 | 4.45 | 4.65 |

**Supplementary table 8.** Total reproduction in 20°C. Post-hoc contrasts.

| Contrast | Estimate | SE | df | t-ratio | p-value |
|---|---|---|---|---|---|
| Control - Fast | 0.06624 | 0.0661 | 25.5 | 1.002 | 0.852 |
| Control - Incr. | 0.09864 | 0.0666 | 26.2 | 1.481 | 0.5832 |
| Control - Slow | 0.06132 | 0.0661 | 25.5 | 0.928 | 0.8836 |
| Control - Warm | 0.05202 | 0.0664 | 25.9 | 0.784 | 0.933 |
| Fast - Incr. | 0.0324 | 0.0656 | 24.7 | 0.494 | 0.9872 |
| Fast - Slow | -0.00493 | 0.065 | 24 | -0.076 | 1 |
| Fast - Warm | -0.01422 | 0.0653 | 24.3 | -0.218 | 0.9995 |
| Incr. - Slow | -0.03732 | 0.0656 | 24.7 | -0.569 | 0.9784 |
| Incr. - Warm | -0.04661 | 0.0659 | 25 | -0.708 | 0.9528 |
| Slow - Warm | -0.00929 | 0.0653 | 24.3 | -0.142 | 0.9999 |
